# Intermediate Migration Yields Optimal Adaptation in Structured, Asexual Populations

**DOI:** 10.1101/003897

**Authors:** Arthur W. Covert, Claus O. Wilke

**Author notes:** Author to whom correspondence should be sent.

## Abstract

Most evolving populations are subdivided into multiple subpopulations connected to each other by varying levels of gene flow. However, how population structure and gene flow (i.e., migration) affect adaptive evolution is not well understood. Here, we studied the impact of migration on asexually reproducing evolving computer programs (digital organisms). We found that digital organisms evolve the highest fitness values at intermediate migration rates, and we tested three hypotheses that could potentially explain this observation: (i) migration promotes passage through fitness valleys, (ii) migration increases genetic variation, and (iii) migration reduces clonal interference through a process called “leapfrogging”. We found that migration had no appreciable effect on the number of fitness valleys crossed and that genetic variation declined monotonously with increasing migration rates, instead of peaking at the optimal migration rate. However, the number of leapfrogging events, in which a superior beneficial mutation emerges on a genetic background that predates the previously best genotype in the population, did peak at the optimal migration rate. We thus conclude that in structured, asexual populations intermediate migration rates allow for optimal exploration of multiple, distinct fitness peaks, and thus yield the highest long-term adaptive success.

## Introduction

The effects of population structure on adaptation have been debated since Sewall Wright first introduced his shifting balance theory, over 80 years ago (Wright 1932, Coyne et al 1997, Wade and Goodnight 1998, Kryazhimskiy et al. 2012). Wright (1932) postulated that populations could only evolve successfully on rugged fitness landscapes if the populations were subdivided into subpopulations or demes, connected by low levels of migration, which could use genetic drift to explore distinct regions of the fitness landscape. Wright theorized that for sexual populations, exploration in structured populations was combined with some form of group selection to drive the overall population to a high fitness peak. Unfortunatly, Wright lacked a useful model system, one with multiple deceptive fitness peaks, to test his theory (Coyne et al 1997, Snigowski and Gerrish 2010, Kryazhimskiy et al. 2012).

How population structure affects adaptation in asexual populations is even less understood. While much work has been devoted to disentangling the factors that determine the speed of adaptation in asexual populations (Muller 1932, Crow and Kimura 1965, Gerrish & Lenski 1998, Papadopoulous 1999, Snigowski and Gerrish 2010, Desai & Fisher 2007, Rouzine et al. 2008, Handel and Rozen 2009), the emphasis in these works has been on large, unstructured populations. Even basic questions about population structure, such as whether it speeds up adaptation or slows it down, and whether this depends on the amount of gene flow and/or the type of fitness landscape, remain unanswered in the asexual case. Some theoretical works suggest that population structure in asexual populations slows down the rate of beneficial substitutions that reach fixation (Gordo and Campos 2006, Pokalyuk et al 2013).

Here, we explore how population subdivision affects adaptation of an asexual population on a rugged fitness landscape. We use self-replicating computer programs (digital organisms) as our model system, and we consider three major elements of asexual adaptation: the role of fitness valleys, the role of genetic diversity, and the role of clonal interference. We find that intermediate migration rates yield the most successful long-term adaptation (i.e., highest fitness). Further, we demonstrate that the benefit of migration consists neither of improved crossing of fitness valleys nor of improved genetic diversity in the population, but rather of a reduction in the effectiveness of clonal interference. At intermediate migration rates, individual clones persist for longer than they would in a large, unstructured population (Habets et al 2007). This increased persistence allows some clones that normally would be destined to go extinct to acquire further beneficial mutations and sweep the population, with a net effect of increased population fitness. In other words: structured asexual populations do many small experiments in local subpopulations before one genotype finally sweeps to fixation, as suggested by Wade and Goodnight (1998).

## Methods

### The Digital Life System Avida

We used Avida version 2.12.8 to simulate the evolution of digital organisms in structured and unstructured environments (Adami 2006, Ofria and Wilke 2004). Digital organisms are computer programs written in a simplified assembly language. An organism's genome is composed of the instructions that make up its program. A standard population of digital organisms is instantiated on a grid of cells, with each cell containing a virtual CPU that can run the organism's genome. A digital organism's main function is to make a copy of itself, and place that copy into one of the adjoining cells in the grid. By default, the cells are connected into a torus, a donut-like shape, so that there are no edges and adjoining cells will always have 8 neighbors into which organisms can place their progeny after self-replication. In our experiments, all populations were seeded with a single ancestor, containing 50 instructions, that could do nothing but self-replicate. All populations where allowed to reach a carrying capacity of 10,000 digital organisms.

Digital organisms self-replicate in three steps. First they must allocate memory to hold their progeny's genome. Second, they must copy each instruction from their code, or their digital genome, into the newly allocated memory. Finally the organisms execute a divide instruction that takes the new copy of their digital genome and places it into an adjoining cell. Digital organisms self-replicate via binary fission, and pass on any mutations in their digital genomes to their progeny. Since digital organisms are computer programs, they would normally replicate perfectly forever. We introduce a small error rate in the divide command that alters a single instruction in the progeny's genome with a given probability. These mutations introduce variation that selection may act on. In all of our experiments we mutated each new digital genome with a 25% chance (one in four offspring organisms experience a mutation), and each mutated offspring differed in only a single point mutation from its parent.

### The Rugged Fitness Landscape of Digital Organisms

The speed at which a digital organism can replicate is determined in part by the speed with which it executes its genome, which is determined by how many instructions its virtual CPU is allowed to execute during each unit of time. During a single update, the unit of time in Avida, all organisms execute on average 30 instructions. However, some organisms get a greater share than others. In particular, digital organisms can increase their CPU speed by computing up to 9 logical functions from binary numbers found in each grid cell (see Lenski et al 2003 for details).

The 9 logical functions range in complexity from relatively simple to relatively difficult (Lenski et al 1999, Lenski et al 2003, Covert et al 2012a, Covert et al, 2013). Organisms that perform more complex logical functions receive a larger share of instruction executions per update than those that execute easier logical functions. In essence, organisms that execute higher-order logical functions have a higher metabolism. However, organisms that are able to execute their instructions more quickly are not guaranteed to have increased fitness. The fittest organism contributes more progeny to the next generation than all other organisms in the population. Since it takes time for digital organisms to self-replicate, organisms must not only get a significant share of instruction executions, but must also use them efficiently to self-replicate as quickly as possible. The trade-off between replication efficiency and the cost of doing complex logical functions drives epistasis in populations of digital organisms (Lenski et al 1999).

### Basic Experimental Protocol

For our initial experiments, we instantiated 7 treatments. 6 of these were structured into 100 subpopulations (or demes) of 100 organisms each and differed only in their migration rate (Figure 1B). Every time a digital organism is born there is a small probability that it will be born into a different deme than its parent's deme. The probability of migration in each treatment differed by an order of magnitude and ranged from 1 migration every 20 births (5×10^−2^) to 1 migration event for every 2,000,000 births (5×10^−7^). The seventh treatment was a single population with an equal total population size of 10,000 organisms. This large single population served as a control for the migration treatments (Figure 1A).

**Figure 1:**
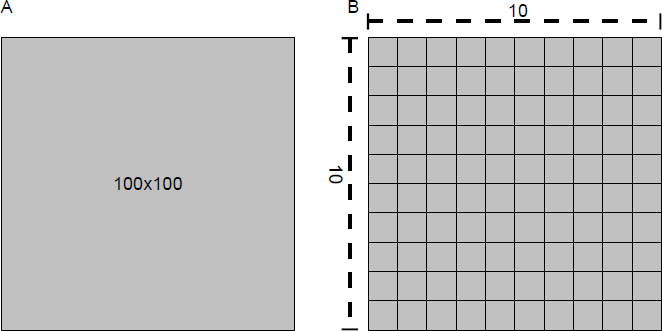
Structured vs. unstructured populations in Avida. Normal Avida experiments (left) take place on an unstructured square grid of 10,000 digital organisms. The grid wraps around itself to form a torus (donut shape) and new digital organisms are born into the adjacent grid cells. In most of our experiments the grid is broken up into 100 sub-populations of 100 digital organisms each (right). After each division, the new digital organism has a small probability of being placed at random into a new subpopulation, resulting in migration.

To insure that each deme was on equal footing, CPU instruction executions were awarded every update to all demes equally, and then awarded to individual organisms within each deme based on the number and type of logical functions they executed. Thus, the metabolism and fitness of a digital organism in one subpopulation in no way impacts digital organisms in neighboring demes, unless there is a migration event. If a highly fit migrant enters a subpopulation with relatively lower fitness, then it will rapidly sweep and fix within that subpopulation. Therefore, we have insured that each structured treatment contains a series of islands, isolated from one another except for rare migration events.

All treatments were instantiated with 50 replicate populations. Experiments were run for 250,000 updates, that allowed for an average of 40,000 generations of evolution (see supplementary materials for a full explanation of generation time effects in our experiments). The fitness of the final dominant genotypes, the most-abundant genotype across all demes at the end of an experiment, was recorded as a measure for the long-term evolutionary success of the population. This experimental setup served as our baseline for testing the effects of population structure.

### Replace Deleterious

Changes in the fitness of digital organisms follow a similar trend to organic study systems: most mutations lower fitness, but some are neutral or even beneficial (Lenski et al 1999). Fitness valleys are created by mutations that are individually deleterious but jointly beneficial (Weinreich et al 2005). Several recent works have found examples of fitness valleys formed by individually deleterious but jointly beneficial mutations in both organic and *in silico* study systems (Covert et al 2013, Cowperthwaite et al 2006, Lenksi et al 2003, Weinreich et al 2006). If such fitness valleys are common enough then they could play an important role in the evolution of structured populations, as originally postulated by Wright (1932).

Experiments with digital organisms have previously shown that passages through fitness valleys can drastically improve the long term adaptive fate of populations (Covert et al 2013). These works have measured the long-term adaptive effects of deleterious mutations by preventing them from entering the population (Covert et al 2013, Covert et al 2012a, and Covert et al 2012b). To prevent deleterious mutations, we proceed as follows (RpD protocol from Covert et al 2013): Whenever a new digital organism is created, it is placed in an isolated test environment to measure its fitness. If its fitness is reduced relative to its parent's fitness, then we revert the deleterious mutation and mutate it again. We repeat this process of reversion and mutation on the organism's genome until we find a mutation that is not deleterious.

By using the replace deleterious (RpD) protocol from Covert et al (2013), we can prevent structured populations of digital organisms from crossing fitness valleys. This replacement method allows us to control for the effects of increased drift in the evolution of structured populations, by observing how well the populations evolve on a rugged landscape when they are unable to cross through fitness valleys. We disabled deleterious mutations in 5 additional treatments to measure the impact of fitness valleys. If structured populations without deleterious mutations have a significantly different distribution of final dominant fitness from our baseline treatments, it would indicate that passage through fitness valleys is a major component of the evolution of structured populations.

### Measuring Diversity in Digital Organisms

Population structure also increases genetic diversity. This increased genetic diversity may be driving the impact of structured populations. Genetic diversity in digital organisms is typically measured by computing the Shannon information entropy on the number of genotypes (Eq. 1). Information entropy is a measure of uncertainty; a higher information entropy indicates a greater uncertainty. For a population of genotypes a higher information entropy indicates that if we draw a random genotype from the population, we have less certainty as to which genotype it will be.

We calculated information entropy as

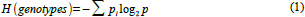

where *p*_*i*_ is the observed frequency of the *i*^th^ genotype. We recorded the information entropy at 200 evenly spaced time points throughout our experiments. We then compared the information entropy at the end of each treatment to the final dominant fitness of that treatment to check for any correlations. If increased diversity is driving the effects of population structure, then we would expect the trend in genetic diversity to be correlated with fitness.

### Leap-frogging in Digital Populations

Finally, we test the theory that population structure increases the probability of leapfrogging. Leapfrogging is a phenomenon that occurs in asexual populations (Gerrish and Lenski 1998, Miller et al 2011). When a mutation gives rise to a beneficial adaptation, its genotype begins to sweep the population, pushing out all other genotypes. Occasionally two genotypes may emerge, with approximately equal fitness. Hypothetically, these two genotypes will sweep out all others, until there are the only two in the population, as shown in Figure 2A (inspired by the work of Muller 1932). We see that genotype c gives rise to two competing genotypes d and e. Genotypes d and e will push the previously dominant genotype, c, out of the population. This sets up a clonal interference scenario: neither genotype has a great enough fitness to overwhelm the other so both will persist until a new beneficial mutation emerges.

**Figure 2:**
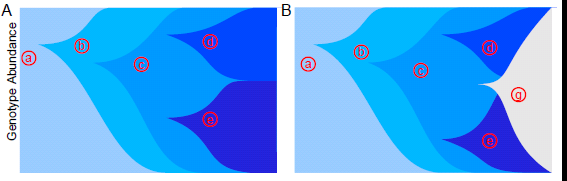
Schematic drawing of clonal interference and leapfrogging. Asexual populations experience repeated selective sweeps. As a new genotype begins to sweep it pushes old genotypes to extinction. These successive selective sweeps are illustrated by genotypes a, b, and c. Each of the early genotypes directly gives rise to its successor, until genotypes d and e emerge at approximately the same time with similar fitnesses. In the top figure, these two genotypes will grow until they push out all other competing genotypes. At that point, genotype e will slowly push genotype d out of the population. The clonal interference depicted in the top figure usually takes a long time to resolve and may slow the overall adaptation of the population. Another scenario, depicted in the bottom figure, involves a third genotype, f, emerging from c after d and e have already begun to sweep the population. In this scenario, f is more fit than both d and e and will rapidly drive both of them extinct. In other words: genotype f leapfrogs genotypes d and e. The defining characteristic of leapfrogging is that a genotype that is dominant at some time (here, genotype e) does not contribute any genetic material to the final population.

Clonal interference may be overcome by a rare mutation that arises before other beneficial mutations overwhelm the population. If a mutation that is clearly superior to either d or e arises, which we label genotype f (figure 2B), it may sweep out genotypes d and e. This scenario is called a “leapfrog” event. In essence, leapfrogging may find the best beneficial mutation from an older starting point, rather than being deceptively drawn to a suboptimal peak.

Beneficial mutations in asexual structured populations may be especially prone to leapfrogging (Woods et al 2011). Any genotype with a beneficial mutation will quickly sweep its own deme, but must wait for a migration event to carry it to other demes, increasing the time it takes to sweep the population (Pokaluk et al 2013). The increased waiting time also increases the probability that a superior genotype will emerge elsewhere in the population before the sweep is complete.

Leapfrogging is extremely difficult to observe in natural study systems, which yield only partial information about which sweeps occurred and when. In digital systems we can record every genotype from the original ancestor to the final dominant genotype; this lineage is called the line of descent. We also recorded the dominant genotype at a series of evenly distributed time points throughout the course of the experiment. We used these data to estimate the rate of leapfrogging by observing how often dominant genotypes are present in the final lineage. If asexual populations exhibit a series of selective sweeps that tend to reach fixation before new mutations can emerge, then every dominant genotype should also be present on the line of descent. Dominant genotypes that are not present on the line of descent have been leapfrogged by a superior genotype. In digital organisms, we can count the number of observed dominant genotypes that are not on the line of descent and use this number as an estimate of leapfrogging.

Our estimation of leapfrogging also includes the possibility that several mutations may accumulate that are epistatically beneficial, though these mutations may not fix in the population. In other words, in our case the leapfrogging genotype could easily represent a combination of several mutations that appeared first in a subpopulation and then swept to fixation together. This scenario is more likely in structured populations, since subpopulations have the chance to explore the fitness landscape independently even if a beneficial mutant exists already elsewhere in another deme. We naively expect any dominant genotype to have the best chance of producing the next dominant genotype. Therefore, our metric assumes that any dominant genotype that is not the line of descent was leapfrogged.

## Results

We analyzed the effect of population structure on rugged landscapes by running experiments across a broad range of migration rates and measuring fitness of the most abundant genotype in each population at the end of the experiment. We ran 6 structured treatments at migration rates ranging from 5×10^−7^ to 5×10^−2^, and a control population without structure. We found that there were significant differences in final dominant fitness among all 7 treatments (*p*<0.001, Kruskal-Wallis test). A rate of 1 migration every 20,000 births (5×10^−5^) produced final dominant fitness significantly higher than the control and all other populations (*p*<0.001, U-test, Table 1 and Figure 3). We refer to this migration rate as the optimal migration rate. A migration rate of 1 migration every 200,000 (5×10^−6^) births also yielded higher fitness than the control population (*p*<0.05, U-test) but still significantly less than the optimal migration rate (both *p*<0.01, U-test). We refer to this migration rate as the suboptimal migration rate. We refer to migration rates between 5×10^−5^ and 5×10^−6^ as ideal migration rates. The range of ideal migration rates extended for at least one order of magnitude around the optimal and suboptimal migration rates.

**Figure 3:**
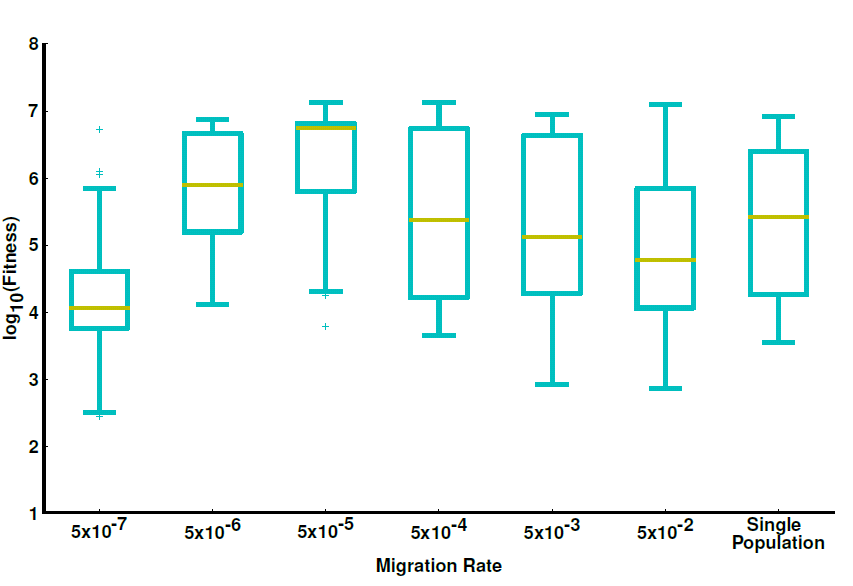
Final dominant fitness vs. migration rate. Each of the boxplots represent the distribution of final dominant fitness in 50 replicate populations. The first six treatments are structured into subpopulations as depicted by Figure 1b. The seventh treatment is an unstructured single population of equal total population size. The median fitness of each treatment forms an inverted v-shaped curve that peaks at the optimal migration rate of 5×10^−5^.

**Table 1:**
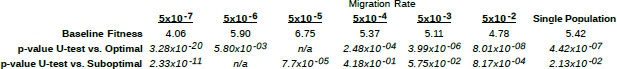
Log_10_ median fitness of all migration rates from the baseline treatments. Fitness values from each treatment were tested against the fitness of the optimal migration rate (5×10^−5^) and the suboptimal migration rate (5×10^−6^). We found that the optimal migration rate yielded significantly higher fitness than all other treatments, and the suboptimal one yielded significantly higher fitness than all other treatments except the optimal one, 5×10^−4^, and 5×10^−3^. Thus, there is a range of ideal migration rates for structured populations, spanning at least one order of magnitude.

We examined the individual differences between treatments to see how different migration rates impacted evolution. Populations with more frequent migration than the optimal did not differ significantly from the control (*p*>0.1, U-test), indicating that structured populations with high rates of migration are qualitatively similar to control populations. The lowest migration rate tested, one migration every 2,000,000 births (5×10^−7^) had significantly lower fitness than all other treatments, including the control (*p*<0.001, U-test). This dip in fitness indicates that certain very low migration rates are similar to having a structured population with a migration rate of 0.0, where sub-populations are too small and too isolated to overcome genetic drift.

To better capture the overall effect of population structure we next plotted only the median final dominant finesses (red line, Figure 4), revealing an inverted v-shaped curve in fitness. The curve peaks at the optimal migration rate and falls off at either end, with higher migration rates being similar to an unstructured population, and lower migration rates being dominated by genetic drift. We tested three different hypotheses on why the optimal migration rate emerged and we used this curve as a baseline for all of our tests.

**Figure 4:**
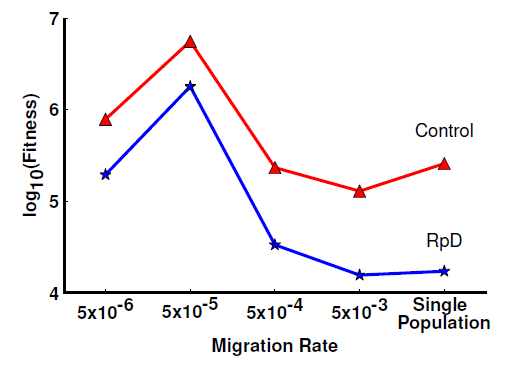
Median final dominant fitness as a function of migration rate, for baseline populations and populations in which deleterious mutations are removed from the population (RpD treatment). The red line corresponds to the baseline treatments (same data as in Figure 3), and the blue line represents the median fitness of RpD treatments. Replacing deleterious mutations significantly reduces fitness in most treatments (Table 2), but does not alter the inverted v-shaped curve that results from varying migration rates. If passage through fitness valleys were driving the effect of population structure, then the effect should collapse when deleterious mutations are turned off, and the blue line should be flat. Instead the effect of population structure is simply shifted down, indicating that whatever effect deleterious mutations are having on the population is independent of the effect of population structure.

### Hypothesis 1: Population structure allows for the traversal of fitness valleys

First, we tested the hypothesis that the effect of structured populations is due to crossing fitness valleys. We instantiated 5 new treatments that replaced deleterious mutations (RpD, see methods for a full explanation). Populations without deleterious mutations are unable to cross fitness valleys. Four treatments consisted of structured populations and migration rates between 10^−6^ and 10^−3^, respectively (one step below the optimal, the optimal, and two steps above), and one additional treatment consisted of a single population control. We compared the median final dominant finesses of each RpD treatment to its corresponding baseline treatment.

We found that while the RpD treatments had significantly reduced fitness relative to most of their controls (see Table 2), they did not remove the inverted v-shaped pattern observed in the base-line treatments (Figure 5). The optimal migration rate was also the significantly higher among most of the RpD treatments (Table 2). If crossing fitness valleys had been the driving force behind the optimal migration rate then we would expect the RpD treatments to form a flat line relative to the unstructured RpD treatment. Instead, we see that the inverted v-shaped curve observed in the baseline treatments is simply shifted down.

**Figure 5:**
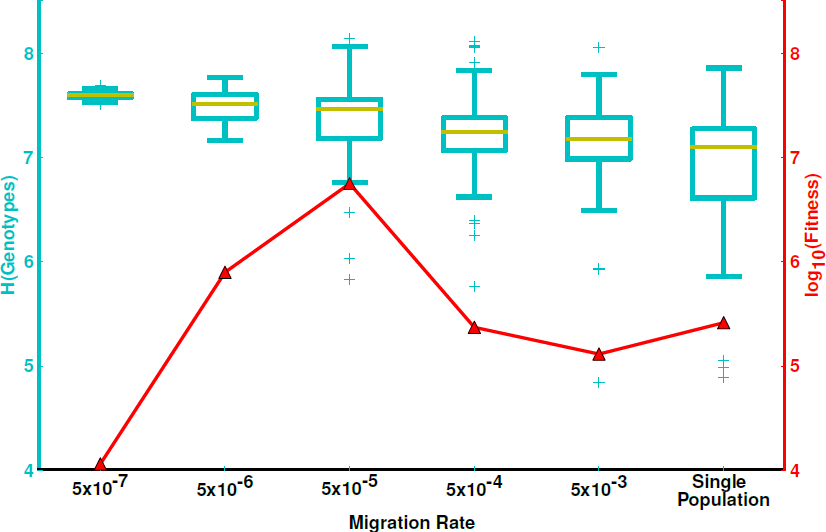
Genetic diversity and final dominant fitness as a function of migration rate. We measure genetic diversity as the information entropy over all genotypes in each final population (box plots, left y-axis), plotted with median fitness of the baseline experiments (red line, right y-axis). Higher information entropy indicates a greater number of genotypes in the final population, and thus more genetic diversity. Genetic diversity is highest at the lowest migration rate and lowest at the highest migration rate. When we compare the trend in genetic diversity to the trend in median final dominant fitness, we see two distinctly different patterns. Genetic diversity is inversely correlated with migration rate, where as final dominant fitness is highest at intermediate migration rates and lower everywhere else. Therefore, while genetic diversity is increased by population structure, it has no meaningful impact on fitness and cannot, by itself, account for the effect of population structure on fitness.

**Table 2:** Log_10_ median fitness and the single population control for the baseline treatments, paired with median final dominant fitness for the replace deleterious treatments (RpD). We found that RpD significantly lowers the overall fitness after adaptation, but fitness is still highest around the optimal migration rate.

|  | Migration Rate |  |  |  | Single Population |
| --- | --- | --- | --- | --- | --- |
|  | <u>5x10<sup>-6</sup></u> | <u>5x10<sup>-5</sup></u> | <u>5x10<sup>-4</sup></u> | <u>5x10<sup>-3</sup></u> |  |
| **Baseline Fitness** | 5.90 | 6.75 | 5.37 | 5.11 | 5.42 |
| **RpD Fitness** | 5.29 | 6.26 | 4.53 | 4.99 | 4.24 |
| **p-value U-test** | n/a | 2.08x10<sup>-03</sup> | 3.23x10<sup>-02</sup> | 1.94x10<sup>-03</sup> | 2.73x10<sup>-02</sup> |

Turning off deleterious mutations did result in lower final fitness relative to the corresponding baseline treatments. This may indicate that traversing a fitness valley is an important event in both structured and unstructured populations. Even if a fitness valley was traversed only once in the evolutionary history of a population, that event may have had a transformative effect on the population's long-term adaptive success (Lenski et al 2003, Cowperthwaite et al 2006, Covert et al 2013). However strong the effect of crossing a fitness valley was, it does not appear to contribute to the effects of asexual population structure.

### Hypothesis 2: Increase in genetic diversity

Next, we measured the genetic diversity of each migration treatment by calculating the genetic entropy (see Methods). Populations with greater diversity will have greater genetic entropy. If increased genetic diversity is driving the effect of structured populations, we expected to see some relationship between genetic entropy and the pattern in final dominant fitness (figure 5). We found that increased genetic diversity was inversely correlated with migration rate. The lowest migration rate (5×10^−7^) had the highest median genetic diversity and the lowest fitness, whereas the optimal migration rate had a mid-range diversity but the highest fitness. The linear trend in genetic entropy makes it unlikely that genetic diversity alone can explain the v-shaped curve caused by population structure. It is possible that some minimal level of increased diversity is necessary for the success of the optimal migration rate, but changes in diversity alone do not tell the whole story.

### Hypothesis 3: Structure yields increased leap-frogging

Finally, we tested the impact of leapfrogging, by measuring the rate of leapfrogging in all of our baseline experiments. We recorded the dominant genotype at 200 equally spaced time points during each experiment, and calculated the percentage of time that the dominant genotype was not present on the line of descent from the final dominant genotype. We excluded the 200^th^ observation from the analysis because it was the final dominant genotype by definition and thus present on the linage 100% of the time. We found that rates of leapfrogging displayed an inverted v-shaped pattern similar to that of fitness, with the highest rate of leapfrogging (97.0%) occurring at the optimal migration rate, and the lowest amount of leapfrogging (88.0%) occurring at the highest migration rate. While differences in the rate of leapfrogging were small in magnitude, they were nevertheless statistically significant. We found significantly elevated leapfrogging at the optimal migration rate relative to all other treatments (*p* < 0.05, Mann-Whitney U-test), except for the suboptimal migration rate (*p* = 0.0558, Mann-Whitney U-test). Thus, elevated rates of leapfrogging appeared to be the main reason why asexual populations had the biggest long-term evolutionary success at intermediate migration rates.

**Figure 6:**
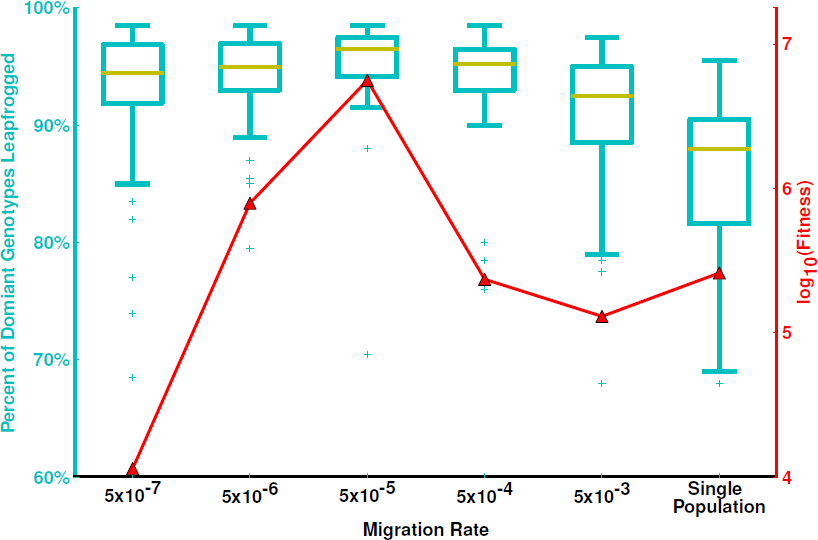
Percent genotypes leapfrogged as a function of migration rate (box plots, left y-axis), plotted with the median fitness of the baseline experiments (red-line, right y-axis). We recorded the dominant genotype at 200 evenly distributed time points during the evolution of each replicate population. For each time point we checked whether the genotype present at that time was an ancestor of the final dominant genotype, and we counted every instance of the dominant genotype not being ancestral to the final dominant as an instant of leap frogging (see methods for a full explanation). We found that there were significantly more instances of leap-frogging at the optimal migration rate than there were at the lowest or highest migration rates, as well as in the single-population control.

## Discussion

We have explored the impact of population structure on the adaptation of asexual digital organisms evolving in a rugged fitness landscape. We have made three key observations: 1. Population structure improves adaptation at an optimal, intermediate migration rate, and it impedes evolution only at very low migration rates. 2. Neither improved crossing of fitness valleys nor an increase in raw genetic diversity fully explain the improvement in adaptation seen in subdivided populations. 3. Instead, the evolutionary success of subdivided populations is highly correlated with the frequency of leapfrogging in these populations. We conclude that intermediate migration rates aid adaptation by creating an increased number of opportunities for leapfrogging events.

While the rate of leapfrogging is significantly increased near the optimal migration rates, the magnitude of the effect is small. This highlights the importance of historical contingency in adaptive evolution. Historically contingent events are rare chance events that have a transformative effect on the long-term adaptation of populations (Blount et al. 2008, Covert et al. 2013). A small number of key mutations, which manage to leapfrog in structured environments at the optimal migration rate but would be unlikely to arise under other conditions, can make a substantial difference in the final fitness that the population can obtain.

Rugged landscapes in particular may require historically contingent mutations (Covert et al 2012a), since these landscapes contain deceptive fitness peaks which confer an immediate fitness gain but limit future adaptation (Woods et al 2011). Structured populations may avoid being trapped on deceptive fitness peaks by leapfrogging around them. Isolated subpopulations can leverage the longer time it takes for a beneficial mutation to sweep the entire population to do more exploration of the fitness landscape. In essence, we show that evolution does many small experiments in structured populations (Wade and Goodnight 1998). Some experiments avoid the deceptive peaks while others do not. Genotypes on deceptive peaks may emerge in structured populations and even dominate for a time, but they have a greater chance of ultimately being swept out by a leapfrogging mutant from another deme that avoided the deceptive peak.

Leapfrogging around deceptive peaks accounts for the importance of population structure on rugged landscapes, but it probably cannot explain what happens on smooth landscapes. Previous work on smooth landscapes (Kryazhimskiy et al. 2012) showed no significant improvement in adaptation from population structure. While some support has been found for rugged (Kivet and Sherlock 2011) and deceptive (Woods et all 2011) landscpaes in populations of simple microorganisms, further examination of the natural world is needed to fully understand the role of fitness landscapes in natural evolution.

Historically it has been argued that the high dimensionality of genotype space will allow populations to follow circuitous paths of neutral mutations to the occasional beneficial adaptation (Van Nimwegen and Crutchfield 2000). While slow, this process of neutral drift and adaptions in large populations has been thought of as the main source of new beneficial mutations. Our work does not directly address how leap-frogging would address this regime of evolution. While the initial stages of digital-organism evolution see a rapid growth in fitness, that growth quickly levels off (as it does in natural populations). Further work is needed to measure the effects of leap-frogging in regimes of smooth, neutral evolution.

Our experiments are similar to past computational experiments done in sexual structured populations to test Wright's Theory of Shifting Balance (Moore and Tonsor 1994). Since our work is asexual Wright's theory does not apply. Furthermore, Wright predicted that the benefit from structured populations would come from increased levels of subpopulation drift that allow for easier passage through fitness valleys. Our experiments explicitly reject the hypothesis that passage through fitness valleys is the driving force behind evolution. Whether similar results would hold for sexual populations is an open question that deserves further research.

In summary, the structure of a population has an important effect on the evolution of asexual organisms on a rugged fitness landscape. At intermediate migration rates, structured populations are able to reduce clonal interference and reach higher levels of fitness than unstructured populations are. Importantly, even though structured populations explore rugged fitness landscapes more effectively, the amount of valley crossing does not appreciably depend on the migration rate. Instead, migration rate seems to affect primarily the extent to which a population can explore multiple fitness peaks at once.

## Acknowledgments

The authors received support from the NSF through the BEACON Center for the Study of Evolution in action (DBI-0939454). Compute time was proved by the Michigan State University High Performance Computing Center and the Texas Advanced Computing Cluster. In addition we thanks Ben Kerr, Charles Ofria, Joshua Nahum, and Jared Carlson-Stevermer. The authors declare that they have no conflicts of interest.

